# Root Ions Fluxes and Osmolarity Changes in Grass Species Differing in Salinity Tolerance

**DOI:** 10.1101/2023.10.16.562608

**Authors:** Liping Wang, Theo Elzenga, Marten Staal

## Abstract

Agricultural areas are increasingly being affected by salt due to irrigation practices and rising levels of salty groundwater. Different plant species have varying degrees of sensitivity to salinity and employ distinct mechanisms to avoid severe damage caused by salt stress. We compared three grass species with different ecological backgrounds, namely *Lolium perenne, Festuca rubra*, and *Puccinellia maritima*, in terms of their ability to maintain growth when exposed to salt stress, the extent of Na^+^-induced K^+^efflux, and the accumulation of salts in their shoots. Our results demonstrate that the changes in K^+^and H^+^fluxes at the root tip induced by NaCl exposure are correlated with the salt tolerance characteristics of these grass species. Specifically, *L. perenne* exhibited the highest leakage of K^+^from its roots, the highest accumulation of Na^+^in its shoots, and the lowest shoot growth under salt stress. On the other hand, *P. maritima* showed minimal changes in ion fluxes in response to salinity stress. *P. maritima* maintained the lowest contribution of Na^+^to the total osmolarity in its shoots and exhibited the least detrimental effect of salt on shoot dry matter. The root cortex including the exodermis and endodermis could be one of the benefit barriers that help defense against salts. In conclusion, root ions fluxes and osmolarity changes in grass species have different salinity tolerance of plants from various habitats. The salt resistance plants restrict leakage of K and exclude Na more effectively. Overall, these results broadened our knowledge of salt resistance in grass species.

## Introduction

Salinity is a major abiotic stress factor that severely limits crop quality, productivity, and poses a threat to food security. Currently, an estima ted 12 million hectares of irrigated land have been rendered unproductive due to salinization, and this area continues to expand rapidly(Nelson & Maredia, 2001; Shrivastava & Kumar, 2015; Chele *et al.*, 2021; Alharbi *et al.*, 2022). Salt stress adversely affects water uptake, leading to physiological drought, and disrupts the ionic balance within plants(Abrol *et al.*, 1988; Hu & Schmidhalter, 2005; Sinclair, 2011; Chele *et al.*, 2021; Singh *et al.*, 2022; Alharbi *et al.*, 2022). Utilizing salt-tolerant crop species and varieties is one potential solution to maintain food productivity and security (Cuartero *et al.*, 2006; Gul *et al.*, 2022; Mousavi *et al.*, 2022; Egea *et al.*, 2023).

Salinity tolerance is a complex trait involving multiple genes and encompasses anatomical, physiological, and biochemical adaptations(Flowers, 2004). Adaptation to salt stress involves osmoregulation, anti-oxidative stress responses, and ion homeostasis(Niu *et al.*, 1995). Salt stress inhibits root growth, prompting the root to undergo changes in cell wall composition, transport processes, cell size and shape, and root architecture(Byrt *et al.*, 2018). The architecture of the root system and the structure of root cells play crucial roles in abiotic stress responses (Uga *et al.*, 2013). Root system architecture (RSA) traits influence nutrient and water uptake under saline conditions(Koevoets *et al.*, 2016; van den Berg *et al.*, 2016; Van Den Berg & Ten Tusscher, 2018). Modifications in cell wall composition, including increased deposition of lignin and suberine, biosynthesis of metalbolism in endodermal and ectodermal cells, help plants maintain water status and alter ion transport characteristics, perform antimicrobial barren against pathogens and predators, thus improve salt tolerance and antimicrobial properties (Shomer *et al.*, 2003; O’Neill *et al.*, 2004; Tocci *et al.*, 2018; Chele *et al.*, 2021; Serra & Geldner, 2022). The binding of Na^+^ ions to cell wall components may impact Na^+^ transfer, affect the binding of other ions, and hinder pectin’s function during cell growth(O’Neill *et al.*, 2004; Byrt *et al.*, 2018). The apoplast, which is involved in intercellular signaling and ion transport during salt stress, is crucial for maintaining high K^+^/Na^+^ ratios(Assaha *et al.*, 2017). In rice, the apoplast barrier has been linked to Na^+^ uptake, and an extensive barrier enhances the plant’s survival under salt stress(Krishnamurthy *et al.*, 2009). In barley, the release of Na^+^ at the root tip appears to occur through apoplastic processes, independent of the salt overly sensitive 1 (SOS1) mediated Na^+^ extrusion as previously hypothesized (Malagoli *et al.*, 2008; Britto & Kronzucker, 2015; Hamam *et al.*, 2016). However, limited information is available on how variation in cell wall structure influences Na^+^ binding and its impact on K^+^ movement.

Maintaining cytosolic K^+^ homeostasis is essential for normal cell functioning and metabolism. Keep ionic balance and preventing Na^+^ (and Cl-) ion toxicity in plant tissues are crucial challenges for salt-stressed plants. Regulation of K^+^ fluxes is a critical process in plant abiotic tolerance(Hasanuzzaman *et al.*, 2019). One adaptive response to high salt exposure is the synthesis of osmotically active small organic compounds and/or antioxidants(Zhu, 2002; Fahad *et al.*, 2015). Vacuolar compartmentalization of Na^+^ through vacuolar Na^+^/H^+^ antiporters prevents the negative effects of Na^+^ in the cytosol and contributes to osmotic potential through Na^+^ accumulation, driving water uptake into the cell

(compartmentalization in the cell wall does not provide the same benefit to maintaining the driving force for water uptake). The investment in osmotic regulation, including ion homeostasis and compartmentalization of Na^+^ (both cytosolic and apoplastic), and the biosynthesis of compatible solutes can negatively affect growth rates. Plants capable of excluding Na^+^ from the shoot potentially maintain high photosynthetic efficiency, rapid growth, and higher yields. Salt-tolerant plants may also exhibit better control of Na^+^ loading into the xylem (Assaha *et al.*, 2017; Wu, 2018). Low Na^+^ accumulation in shoot tissue is often used as a robust indicator for screening salinity tolerance(Møller *et al.*, 2009). However, a study on barley suggested that the ability of plant cells to retain K^+^, rather than restricting Na^+^ uptake, may be the critical trait for salt tolerance(Chen *et al.*, 2005).

Varieties of quinoa (Chenopodium quinoa Wild) from high saline natural habitats exhibited reduced stomatal density and increased leaf sap K^+^ concentrations(Shabala *et al.*, 2013).. In moderately salt-resistant wheat leaves, K^+^ uptake significantly decreased under salt stress(Hu & Schmidhalter, 2005). Maintaining a high cytosolic K^+^/Na^+^ ratio, whether through restricting Na^+^ accumulation or preventing K^+^ loss, is a crucial determinant of plant salt tolerance (Nublat *et al.*, 2001; Tester & Davenport, 2003; Colmer *et al.*, 2006; Shabala & Cuin, 2008; Barrett-Lennard & Shabala, 2013; Anschütz *et al.*, 2014). A strong correlation has been observed between the ability of plant roots to restrict Na^+^-induced K^+^ efflux and plant salinity tolerance(Chen *et al.*, 2015). The membrane potential, regulated by plasma membrane (PM) H^+^-ATPase activity, is one of the key factors governing Na^+^-induced K^+^ efflux(Chen *et al.*, 2007b; Sun *et al.*, 2009). Salt-resistant barley cultivars exhibited increased H^+^-ATPase activity and a more hyperpolarized membrane potential, effectively preventing Na^+^-induced K^+^ efflux (Chen *et al.*, 2007b,a).

In this study, we employed the Microelectrode Ion Flux Estimation (MIFE) technique to measure osmotic adjustment and ionic flux characteristics along the root surfaces of three grass species: *Lolium perenne, Festuca rubra*, and *Puccinellia maritima*. These perennial grasses have distinct ecological backgrounds. *Lolium perenne* is a common grassland species used for fodder and biofuel (Jauhar, 1993; Newman, 2001; Kunkel *et al.*, 2006; Wang & Ge, 2006). *Festuca rubra* and *Puccinellia maritima* are dominant species in salt marshes(Rouger & Jump, 2015). *Festuca rubra* has a broad ecological range(ASHRAF *et al.*, 1986), while *Puccinellia maritima* is limited to saline areas such as coastal salt marshes. *Puccinellia maritima* is favored as a food source for geese and hares in salt marshes due to its ability to prevent Na^+^ accumulation in leaves while adjusting osmotically through the synthesis of small organic osmolytes, making it more palatable(Fokkema *et al.*, 2016). We investigated the effects of salt exposure on root ion fluxes in these three grass species, considering their varying characteristics in coping with high NaCl levels.

## 2. Materials and Methods

### 2.1. Plant material

Three species, namely *Lolium perenne, Festuca rubra*, and *Puccinellia maritima*, were selected to determine ion flux characteristics. Seeds of *L. perenne* and *F. rubra* were obtained from Samenshop24 in Früchte Röben, Germany. Seeds of *Puccinellia maritima* (Plant ID: PI 260702) were provided by the U.S. National Plant Germplasm System, collected from saline habitats in Afghanistan.

### 2.2. Leaf sap osmolarity measurements

To study leaf osmolarity responses to salinity, experiments were conducted in a climate chamber (21°C, RH 50%, 16/8h day/night cycle, light intensity of 100 ± 20 μmol m−2 s−1). Seeds of each species were sown directly into plastic pots filled with vermiculite. After seven days of germination, seedlings were thinned to 20 equal-sized individuals. The plants were irrigated daily with a 25% strength Hoagland solution. Salt treatments began three weeks after germination, with daily increments of 25 mM NaCl until reaching the final treatment levels: 0 mM (control), 50 mM, 100 mM, 175 mM, and 225 mM NaCl (n=3). After seven days of exposure to salinity, leaf sap was collected by squeezing the tissue using a garlic press(Zaki Mostafa Ali, 2017). The osmolarity of the leaf sap was determined using a Wescor HR-33T dew point microvoltmeter. The electronic conductivity (EC) of the leaf sap was measured using a LAQUATwin (model S070, MFG No: EM5K0529). The EC values were used to calculate the contribution of NaCl to the total osmolarity. Three replicates per treatment were measured, with each sample having three measuring replicates.

### 2.3. Seedling preparing for Ion flux measurement

Seeds were sown on sterilized filter paper in square Petri dishes (12x12x1.5 cm3), with ten seeds per dish and three replicate dishes for each species. The Petri dishes were placed in the climate chamber described earlier and watered daily with distilled water. Two-day-old seedlings were transferred to 30 L containers filled with a 25% strength Hoagland solution. Ion flux measurements were conducted using six-day-old seedlings.

### 2.4. Ion-specific micro-electrodes

Microelectrodes were prepared and silanized following the method described by Zaki Mostafa Ali(Zaki Mostafa Ali, 2017). Further details on electrode preparation and calibration can be found in Box S1 and S2 in the supplementary materials (SM). H^+^-selective microelectrodes were back-filled with 15 mM NaCl and 40 mM KH2PO4, with the electrode tips front-filled with Hydrogen Ionophore I-cocktail A (Fluka 95297). K^+^ electrodes were back-filled with a 200 mM KCl solution and front-filled with Potassium Ionophore I-Cocktail A (Fluka 60031). The electrodes were calibrated using standard solutions, and only electrodes meeting specific criteria were used in the experiments.

### 2.5. Flux measurement

The root of a six-day-old seedling was placed in a small Petri dish (5 cm diameter) on an inverted microscope (Nikon TMS, Tokyo, Japan) during the measurements. The Petri dishes were filled with 5 ml of MIFE Bath Solution (BS), which contained 200 μM MgCl2, 100 μM CaCl2, 100 μM KCl, and 0.5 mM MES (pH 6.0), supplemented with the required NaCl concentration for the salt treatment.

NaCl-induced ion fluxes along the root’s distal section were determined by incubating the seedlings for 30 minutes in a salt solution(Zaki Mostafa Ali, 2017) prior to ion flux measurements. Three salt treatments were applied: 0 mM NaCl (control), 100 mM NaCl, and 200 mM NaCl in BS. Ion fluxes were recorded along the surface of the primary root at specific distances from the root tip. Measurements were taken at 0.25, 0.50, 0.75, 1.00, 1.25, 1.50, 2.00, 2.50, 3.00, 3.50, 5.50, and 7.50 mm from the root tip, and each flux measurement was averaged over a 2-minute duration. H^+^ and K^+^ fluxes were measured separately.

Instantaneous changes in ion fluxes induced by NaCl addition in the elongation zone of the primary root (approximately 2.00 mm from the root tip) were determined. Ion fluxes were measured in BS without salt for around 10 minutes to stabilize. Then, a final concentration of 50 mM NaCl was added. After approximately 25 minutes, an additional 50 mM NaCl was added (totaling 100 mM NaCl), and fluxes were recorded for another 15-25 minutes.

### 2.6. Suberin staining

To observe suberization of epidermal and endodermal cell walls, root tips measuring 5 cm in length were embedded in 6% agarose. Cross-sections of the embedded roots were made using a hand microtome (MT-5503, Euromex). The sections, with a thickness between 50 and 100 μm, were incubated overnight at room temperature in 1% (w/v) berberine hemi-sulfate (Sigma). After rinsing the sections with distilled water, excess stain was removed. Root sections were observed using epifluorescence on a Zeiss Axiophot microscope with a UV filter set (excitation filter BP 365, dichroic mirror FT 395, barrier filter LP 397; Zeiss, Oberkochen, Germany).

### 2.7. Data analysis

Data analysis and graphing were performed using Prism 9.0 (GraphPad Software, San Diego, USA). Two-way ANOVA analysis was conducted on osmolarity data, with a significance level of P<0.05. Three-way ANOVA analysis was performed using SPSS 28.0.1.0 (IBM, Armonk, USA).

## 3. Results

### 3.1. Differences in salinity sensitivity between the three grass species

The three grass species selected for the study exhibited different levels of salinity tolerance. *Lolium perenne*, considered moderately salinity tolerant, showed greater shoot growth reduction compared to *Festuca rubra* and *Puccinellia maritima* when exposed to 50 mM and 100 mM NaCl in the growth medium. The difference in salinity tolerance are evident from Figure S2 and Table S3 in SM, showing that shoot growth (expressed as dry weight) of *P. maritima* is less affected than the other two species by 50 and 100 mM NaCl in the growth medium.

### 3.2. Ion flux profiles along the roots are species specific and affected by NaCl

Ion flux profiles along the distal section of the primary root varied among species and were influenced by the presence of NaCl in the medium (Figure 1) . Under control conditions, only a small efflux of H^+^ and K^+^ ions was observed along the root, with some variations between species. Upon incubation in NaCl for 30 min, the K^+^ efflux increased and exhibited a more pronounced pattern, including an expanded peak zone, higher peak efflux, and slight efflux in the proximal zone of the root (Figure 1B, D, F. Table S1 in the SM). The H^+^ flux profile differed among species, with *L. perenne* showing a change from H^+^ influx to efflux in the presence of NaCl (Figure 1A), *F. rubra* displaying minimal changes (figure 1C), and *P. maritima* already exhibiting a considerable H^+^ efflux even without NaCl, which was further increased by NaCl exposure (Figure 1E).

**Figure 1.**
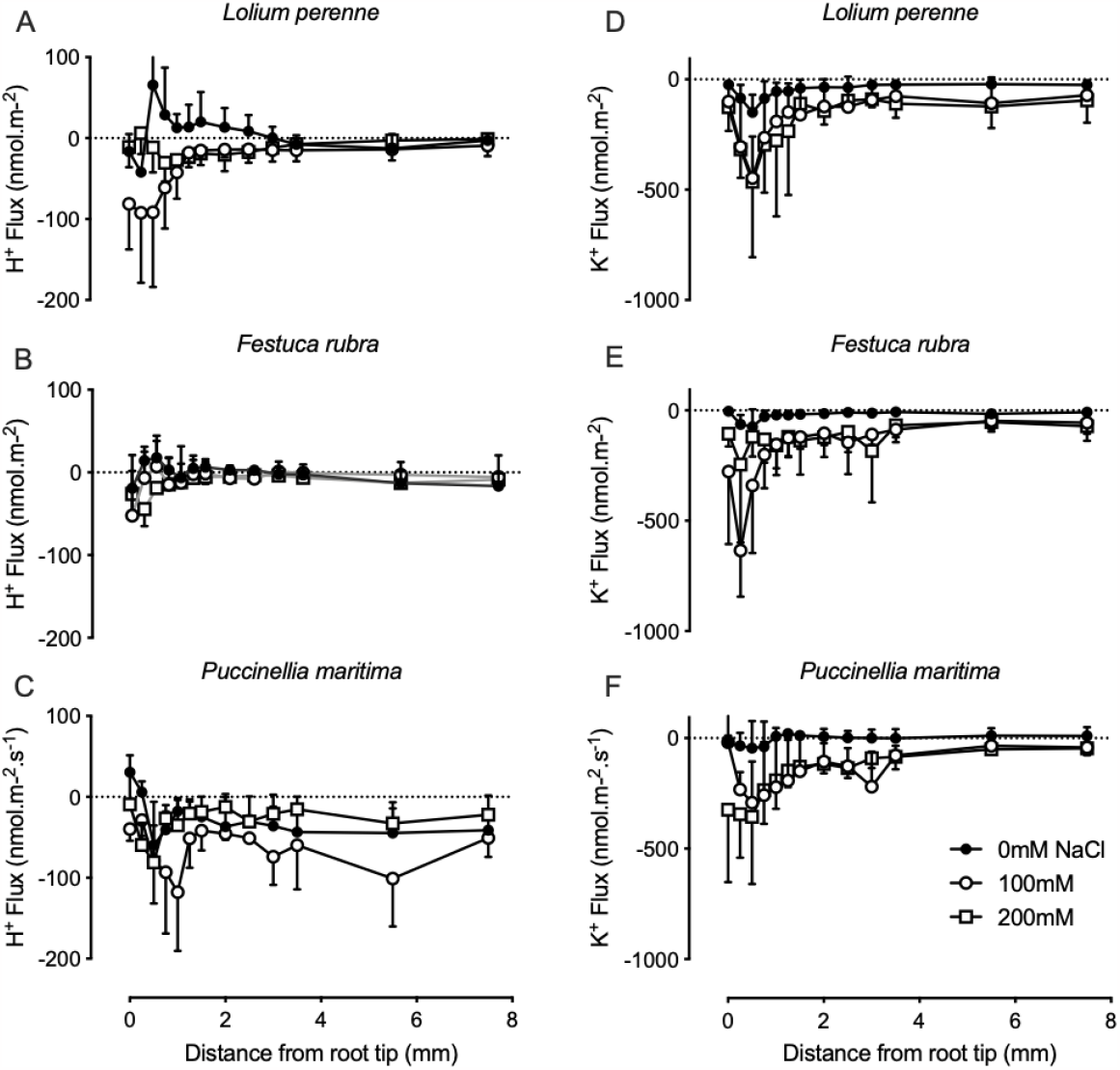
The net H^+^and K^+^ion fluxes along the roots of *Lolium perenne, Festuca rubra*, and *Puccinellia maritima* were measured using ion-selective electrodes. The H^+^flux along the root of each species is shown in the left column (A, B, and C for *Lolium perenne, Festuca rubra*, and *Puccinellia maritima*, respectively), while the K^+^flux is shown in the right column (D, E, and F for the three species, respectively). The roots were measured under three salinity levels: 0 mM NaCl (control), 100 mM NaCl, and 200 mM NaCl. The number of seedling replicates ranged from 3 to 7.A three-way analysis of variance (ANOVA) was performed using SPSS (version 28) with species (*P. maritima, L. perenne, and F. rubra*), salt concentration (0, 100, and 200 mM NaCl), and root zone (0-1 mm, 1-2 mm, and >2 mm) as independent variables. The data used for the analysis can be found in the Supplemental Material (SM TableS4.1 and S4.2). The analysis revealed significant effects of species (P<0.001), salt treatment (P<0.01), and root zone (P<0.05) on the H^+^flux. There were also significant interactions between salt treatment*species and salt treatment*root zone for the H^+^flux (P<0.01).Regarding the K^+^flux, the analysis showed significant effects of salt treatment and root zone (both P<0.001). There was also a significant interaction between salt treatment and root zone for the K^+^flux (P<0.05). These results indicate that species, salt treatment, and root zone all have significant effects on both H^+^and K^+^fluxes, with interactions between salt treatment and species, as well as salt treatment and root zone, being particularly significant for the H^+^flux. The K^+^flux is influenced by salt treatment and root zone, with a significant interaction between salt treatment and root zone.

### 3.3. Instantaneous NaCl-induced H^+^ and K^+^ flux changes

Roots exposed to a sudden increase of NaCl in the external medium respond with an instantaneous change in H^+^ fluxes in a distinct, species-specific way (Figure 2A, C and E). Before the addition of NaCl, *L. perenne* showed a small H^+^ influx, while *F. rubra* and *P. maritima* exhibited a small efflux, on average 58, 12 and 34 nmol·m^-2^·s^-1^, respectively.

**Figure 2.**
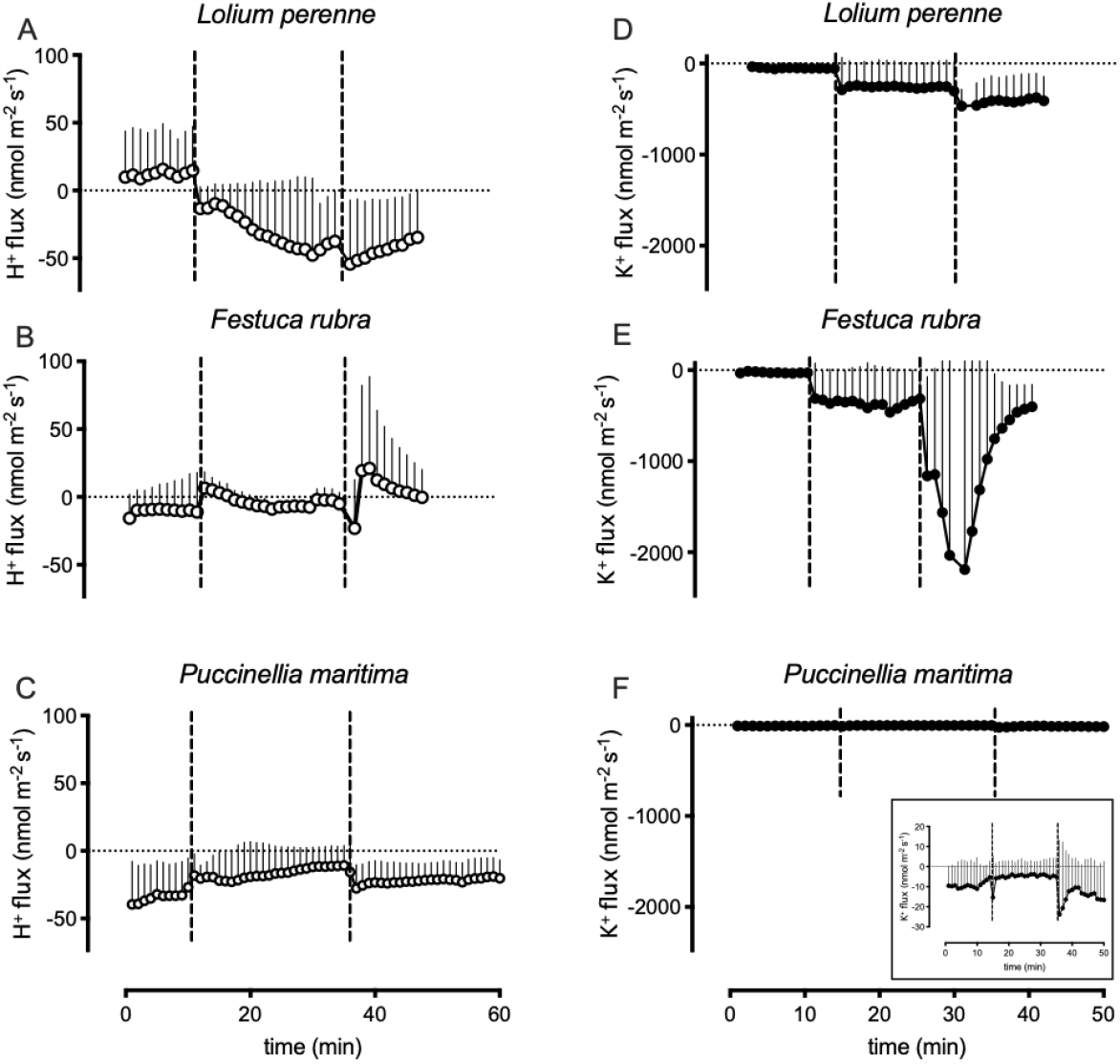
The instantaneous changes in net H^+^and K^+^fluxes induced by salt addition in *Lolium perenne, Festuca rubra*, and *Puccinellia maritima*. The kinetic H^+^flux of the root for each species is depicted in plots A, B, and C, while the kinetic K^+^flux is shown in plots D, E, and F, respectively. The roots were placed in a 5 ml MIFE Bath Solution containing 200 μM MgCl2, 100 μM CaCl2, 100 μM KCl, and 0.5 mM MES (pH 6.0). The fluxes were measured for 10 minutes, after which 5 ml of 100 mM NaCl MIFE BS was added to bring the NaCl concentration to 50 mM, and the fluxes were measured for an additional 15-25 minutes. Finally, another 5 ml of 200 mM NaCl MIFE BS was added, resulting in a final NaCl concentration of 100 mM, and the measurement continued for another 15 minutes. The average of 10-second recordings was taken for each data point, with replicates ranging from 6 to 8. A three-way ANOVA was performed using SPSS (version 28) with species (*P. maritima, L. perenne,* and *F. rubra*), salt concentration (0, 50, and 100 mM NaCl), and time period after the addition of the salt solution (early, middle, and late) as independent variables. The data used for the analysis can be found in the Supplemental Material (SM Table S5.1 and S5.2). The analysis revealed a significant effect of species (P<0.001) on both H^+^and K^+^fluxes. There was also a significant interaction between salt treatmentspecies for the H^+^flux (P<0.001). In the case of the K^+^flux, there were significant effects of species (P<0.001) and salt treatment (P<0.01), as well as a significant interaction between speciessalt treatment (P<0.05). These results indicate that species has a significant effect on both H^+^and K^+^fluxes, with specific interactions between salt treatment and species for the H^+^flux. The K^+^flux is influenced by both species and salt treatment, with an additional interaction between species and salt treatment

**Figure 3.**
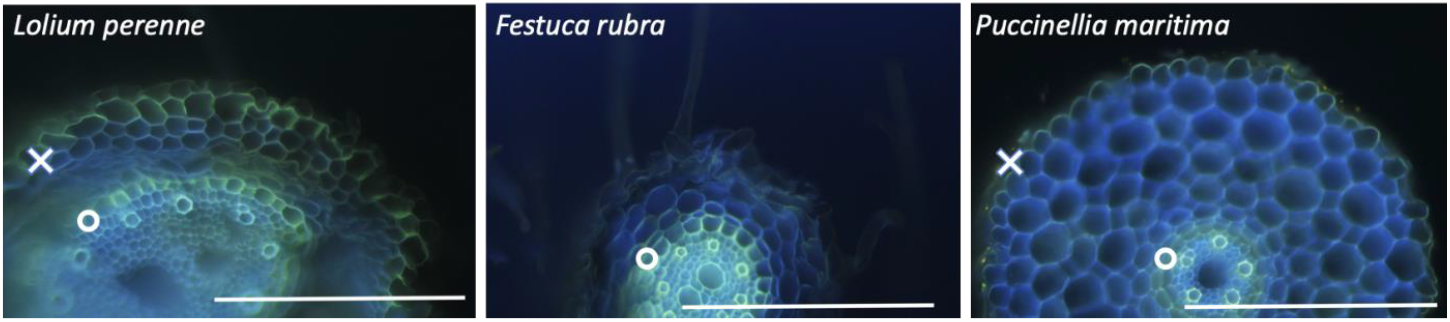
Epifluorescent images of root cross sections stained with berberine visualizing the suberization of the cell walls of *Loloim perenne, Festuca rubra* and *Puccinellia maritima*. The greenish fluorescence observed is a result of suberine staining by berberine, while the whitish fluorescence represents the autofluorescence of vessel elements and tracheids in the xylem. The clear blue autofluorescence corresponds to the cell walls. In the figures, the ‘X’ indicates the location of the exodermis, which is absent in *F. rubra*, while the ‘O’ represents the location of the endodermis. These figures are representative examples of root cross sections from five individual plants of each species. In all five samples of *F. rubra*, only the endodermis was stained, indicating the absence of the exodermis. On the other hand, in all five samples of *L. perenne* and *P. maritima*, suberine staining was observed in both the endodermis and the exodermis. These findings demonstrate that there are anatomical differences in suberization between the three grass species, with *F. rubra* lacking the exodermis layer observed in *L. perenne* and *P. maritima*.

When NaCl was added to the external medium, the three species responded with distinct changes in H^+^ and K^+^ fluxes (Figure 2*). L. perenne* exhibited a sharp increase in H^+^ efflux upon 50 mM NaCl addition, which gradually increased over 35 minutes. A further increase in NaCl concentration in the medium to 100 mM did not further increase the H^+^ efflux but rather a slow decline. In contrast, *F. rubra* showed a small change in H^+^ flux from efflux to influx upon 50 mM NaCl addition, followed by another small influx upon 100 mM NaCl addition. *P. maritima* displayed minimal changes in H^+^ flux in response to NaCl.

Regarding K^+^ flux, *L. perenne* and *F. rubra* exhibited stepwise increases in efflux upon NaCl addition, with *F. rubra* displaying a transient large increase before returning to the original level in about 10 min. *P. maritima* showed minimal changes in K^+^ efflux compared to the other two species. On average, the efflux of K^+^ exposed to 50 mM NaCl was just -5 nmol·m-2·s-1, only a fraction of the efflux observed in *L. perenne* and *F. rubra*. Increasing the NaCl to 100 mM NaCl slightly changed the K^+^ efflux to an initial maximum of about -25 nmol·m-2·s-1.

### 3.4. Anatomical differences in the young root tips

Cross sections of the roots revealed anatomical differences between the species. Both *L. perenne* and *P. maritima* displayed a suberized exodermis alongside a strongly stained endodermis. In contrast, *F. rubra* only showed staining of the endodermis, with no clear presence of an exodermis.

### 3.5. NaCl contribution to leaf sap osmotic potential

The total osmolarity of leaf sap differed among species and salinity levels. All species increased their total osmolarity under NaCl treatment compared to the control (Figure 4). L. perenne showed the most significant increase in osmolarity. Under control conditions, *L. perenne* had an osmolarity of 241 mOsm, *F. rubra* 233 mOsm, and *P. maritima* 206 mOsm. The total osmolarity of *L. perenne* increased from 514 mOsm to 561 mOsm when NaCl concentration increased from 50mM to 100mM, and further to 912 mOsm and 1224 mOsm in 175 mM and 225 mM NaCl, respectively. Compared with *L. perenne, F. rubra* and *P. maritima* increased much less, only 279 mOsm and 309 mOsm in 100mM NaCl and 372 and 587 mOsm in 225mM NaCl, respectively.

**Figure 4.**
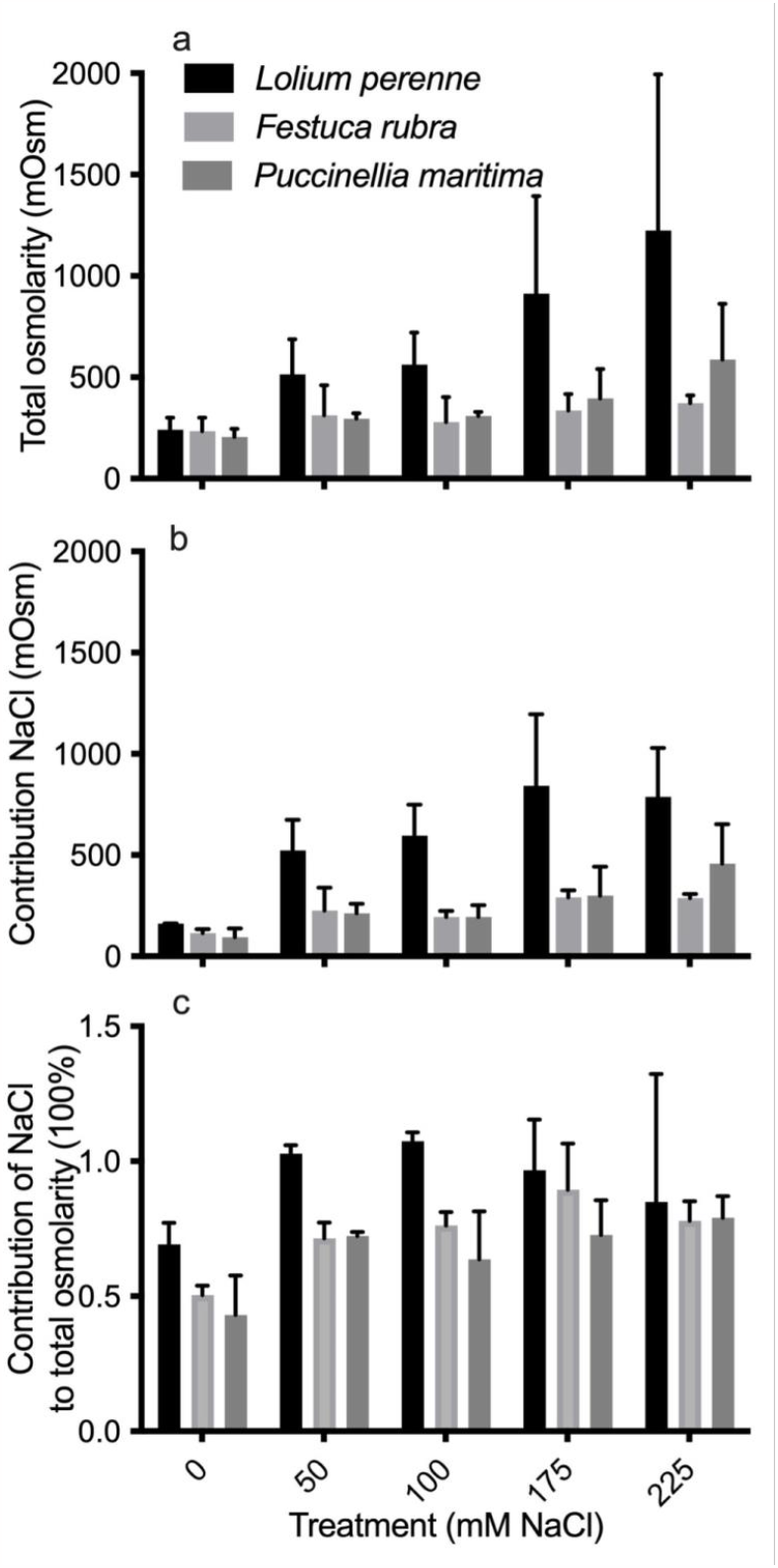
Leaf sap osmolarity and the contribution of NaCl to the total osmolarity of leaf sap of *L. perenne, F. rubra* and *P. maritima*. The plants were grown in pots filled with vermiculite and received daily watering with a 25% Hoagland solution. Stepwise NaCl treatments were applied at concentrations of 0 mM (control), 50 mM, 100 mM, 175 mM, and 225 mM. The above-ground leaves of the plants were collected, and the leaf sap was extracted by squeezing them in a garlic press. The osmotic potential and electric conductivity of the leaf sap were measured to determine the total osmolarity and the percentage contribution of NaCl to the total osmolarity. (a) Total osmolarity of leaf sap: The graph shows the total osmolarity of the leaf sap in the three species. Significant differences were observed between the different salt treatments, indicating that the osmolarity of the leaf sap increased with increasing NaCl concentration. (b) NaCl contribution to osmolarity: This graph illustrates the contribution of NaCl to the total osmolarity of the leaf sap. It shows that as the NaCl concentration increased, the contribution of NaCl to the osmolarity also increased. Significant differences were found between the different salt treatments, indicating that NaCl made a larger contribution to the osmolarity at higher NaCl concentrations. (c) Percentage of NaCl in total osmolarity: This graph represents the percentage of NaCl in the total osmolarity of the leaf sap. It shows that the percentage of NaCl contribution to the total osmolarity varied among the species and salt treatments. No significant interactions were observed between the salt level and species in terms of osmolarity and the percentage of NaCl contribution to osmolarity. However, significant differences were found between the different salt treatments and species. Overall, these results demonstrate that the leaf sap osmolarity increased with NaCl treatment, and NaCl made a significant contribution to the total osmolarity in the three grass species.

The contribution of salt to the total osmolarity in salt-treated plants increased compared with control plants (P<0.05, Figure 4 B, C). *L. perenne* showed the highest increase, with the percentage of NaCl contribution to total osmolarity being the highest among the three species at all salt concentrations. *F. rubra* and *P. maritima* exhibited lower increases in osmolarity and lower percentages of NaCl contribution compared to *L. perenne*.

## 4. Discussion

### 4.1. Root ion fluxes and salt tolerance in grass

The overall profiles of K^+^ fluxes along the root tip and expansion zone in the presence and absence of NaCl resemble those described in other species(Shabala & Cuin, 2008). The fluxes start from a net zero at the ultimate tip of the root and increase to a maximum at around 1 to 2 mm behind the tip. Subsequently, they gradually decrease to very low values through the expansion zone

Earlier papers described the relationship between Na^+^-induced K^+^ fluxes and salinity tolerance in maize(Zaki Mostafa Ali, 2017) and barley(Chen *et al.*, 2005). In those studies, it was assumed that species capable of minimizing K^+^ loss from the root cortical cells could also prevent the transport of Na^+^ into the central cylinder and ultimately prevent Na^+^ accumulation in the leaves.

In Figure 5, we present a hypothetical model that outlines the relevant transport processes occurring in cortical cells following exposure to Na^+^. Although molecular data specific to the three grass species in this study are not available, the model is based on studies conducted on other species. The model depicts a cascade of interrelated processes that are triggered by the stepwise increase in external Na^+^ concentration. According to the model, positive charges carried by Na^+^ enter root cells through non-selective cation channels (NSCC). This influx of Na^+^, along with the activity of HKT-type low-affinity transporters and possibly K^+^-selective ion channels like AKT1, can lead to the accumulation of Na^+^ in the cell, causing depolarization of the membrane potential. The depolarization of the membrane induces K^+^ efflux and activates H^+^-pumping ATPase, resulting in H^+^ efflux. The combined efflux of K^+^ and H^+^ can re-hyperpolarize the membrane potential, potentially reversing the K^+^ flux from efflux to influx. The H^+^ efflux through H^+^-pumping ATPase also increases the proton motive force, which is the force across the membrane generated by the combined electrical potential and pH differences. This increased proton motive force can facilitate the efflux of Na^+^ and the active uptake of K^+^. The model suggests that second messengers such as Ca2^+^, cGMP, and reactive oxygen species (ROS) in the cytoplasm may further influence K^+^ uptake and downstream transcription of genes involved in salinity responses It’s important to note that in this study, only the K^+^ and H^+^ fluxes were measured, and their sizes are expected to be proportional to the Na^+^ flux into the root cortical cells based on the model.

**Figure 5.**
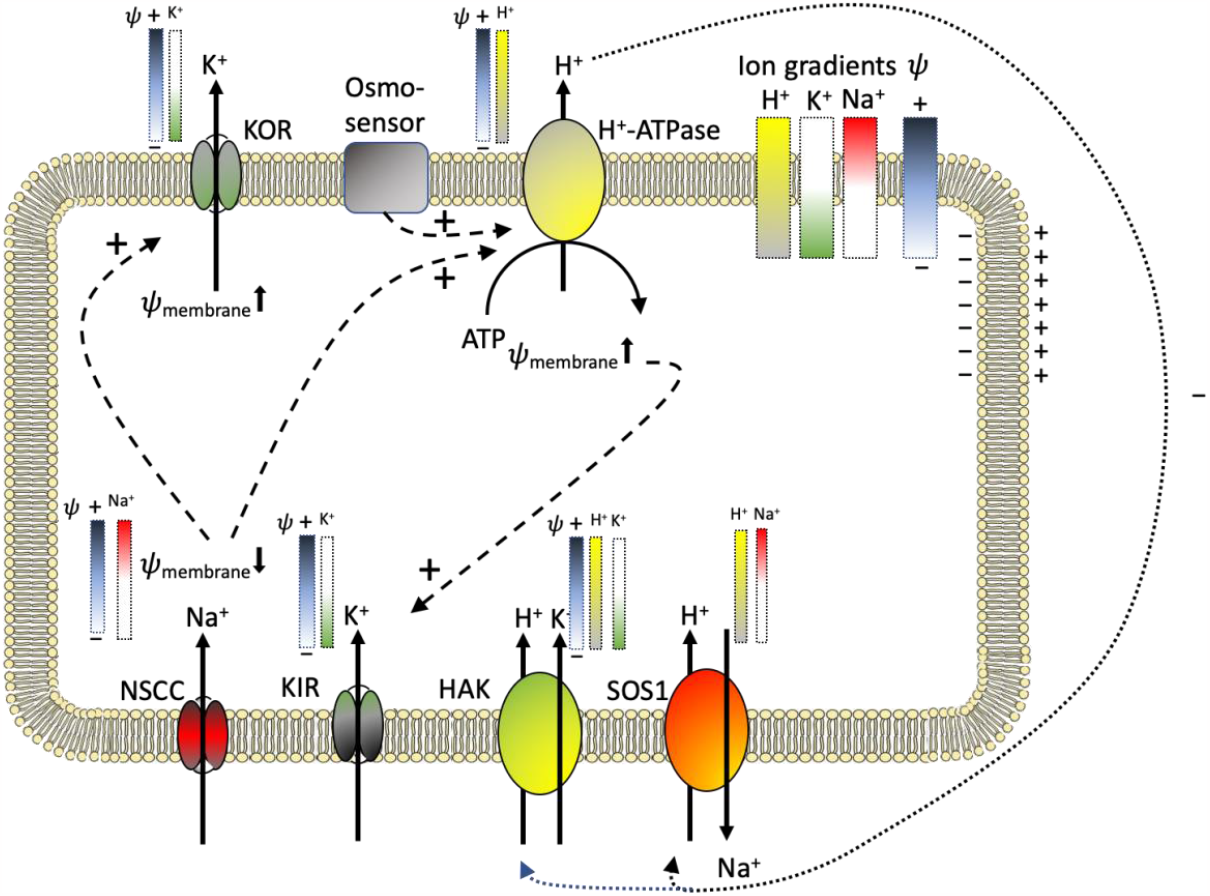
Hypothetical model, based on studies on species of which the molecular data on transporters is known, of the membrane transporters mediating and the forces acting on the fluxes of H^+^, K^+^ and Na^+^ involved in the responses of root cortex cells when challenged with increased external Na^+^ concentrations. In the model, the transporter proteins are represented by symbols, and the arrows indicate the direction of flux through these transporters. Positive interactions between components are indicated by dashed arrows labeled with a ‘^+^’ sign. The gradient bars next to the transporters represent the different components, such as the membrane potential and ion concentration differences across the membrane, that contribute to the forces acting on the fluxes. Abbreviations used in the model include KOR (K^+^ outward rectifying channel), KIR (K^+^ inward rectifying channel), H^+^-ATPase (proton-pumping ATPase), HAK (high-affinity K^+^ transporter, also known as HUK/HAK/KT, a K^+^/H^+^ symporter), NSCC (non-selective cation channel), and SOS1 (salt overly sensitive 1, a Na^+^/H^+^ antiporter). The sequence of events is assumed to be as follows: positive charges carried by Na^+^ enter root cells through non-selective cation channels (NSCC) (Amtmann & Sanders, 1998). The Na^+^ influx, facilitated by HKT-type low-affinity transporters and possibly K^+^-selective ion channels like AKT1, can lead to significant accumulation of Na^+^ within the cell(Yuan *et al.*, 2005), resulting in the depolarization of the membrane potential. The depolarization of the membrane potential induces K^+^ efflux, as well as activates H^+^-pumping ATPase, which leads to H^+^ efflux. The combined efflux of K^+^ and H^+^ ions helps re-hyperpolarize the membrane potential, potentially reversing the K^+^ flux from efflux to influx. The efflux of H^+^ ions through H^+^-pumping ATPase increases the proton motive force, which is the force across the membrane generated by the combined electrical potential and pH differences. This proton motive force can power the efflux of Na^+^ and the active uptake of K^+^. The model suggests that second messengers such as Ca2^+^, cGMP, and reactive oxygen species (ROS) present in the cytoplasm, which are involved in salinity signaling, may subsequently promote K^+^ uptake and affect downstream transcription of numerous genes(Kiegle *et al.*, 2000; Donaldson *et al.*, 2004; Yuan *et al.*, 2005).

When exposed to high external NaCl, the differences in ion flux characteristics at the root surface among the three species may be correlated with their variations in salinity tolerance and root cell composition. *L. perenne* and *P. maritima* exhibit suberization of the exodermis and intensely stained endodermis, suggesting that these plants prevent NaCl uptake. On the other hand, *F. rubra* lacks suberization in the exodermis, which may contribute to higher K^+^ leakage caused by NaCl entry. Suberinization of the cell walls in both the endodermis and exodermis can influence the regulation of solute transport(Enstone *et al.*, 2002; Barberon, 2017). In rice (Oryza sativa), exposure to salt increases suberin deposition in the roots, resulting in reduced Na^+^ accumulation in the shoots(Krishnamurthy *et al.*, 2009).

Although the flux measurements provide quantitative data on K^+^ and H^+^ fluxes only, they also offer a semi-quantitative estimate of Na^+^ fluxes. It is assumed that the initial K^+^ efflux is triggered by the depolarization caused by Na^+^ influx and is at least partially correlated quantitatively. Following this assumption, the species with low or zero K^+^ efflux, indicating minimal depolarization, are expected to have higher Na^+^ tolerance(Shabala & Cuin, 2008). In the current study, *P. maritima*, the species with the highest salt tolerance, exhibits the lowest Na^+^-induced K^+^ efflux and shows rapid recovery after the initial flux change. A report on P. tenuiflora, a closely related halophytic grass, highlights the role of SOS1, HKT1;5, and NHX1 in Na^+^ homeostasis and provides evidence that regulating salt sequestration in the roots limits Na^+^ accumulation in the shoots(Zhang *et al.*, 2013).

The data also reveal a relationship between K^+^ efflux characteristics and Na^+^ accumulation in the shoot, as depicted in Figure 6. Additionally, the peak K^+^ efflux, steady-state K^+^ efflux, and The (lack of) recovery K^+^ efflux are associated with the total Na^+^ accumulation in the shoot(data not shown). Figures 6 suggests that the integrated K^+^ efflux characteristics are related to the total Na^+^ accumulation in the shoot and can be indicative of the species’ response to salinity(Ashraf et al., 2023).

**Figure 6.**
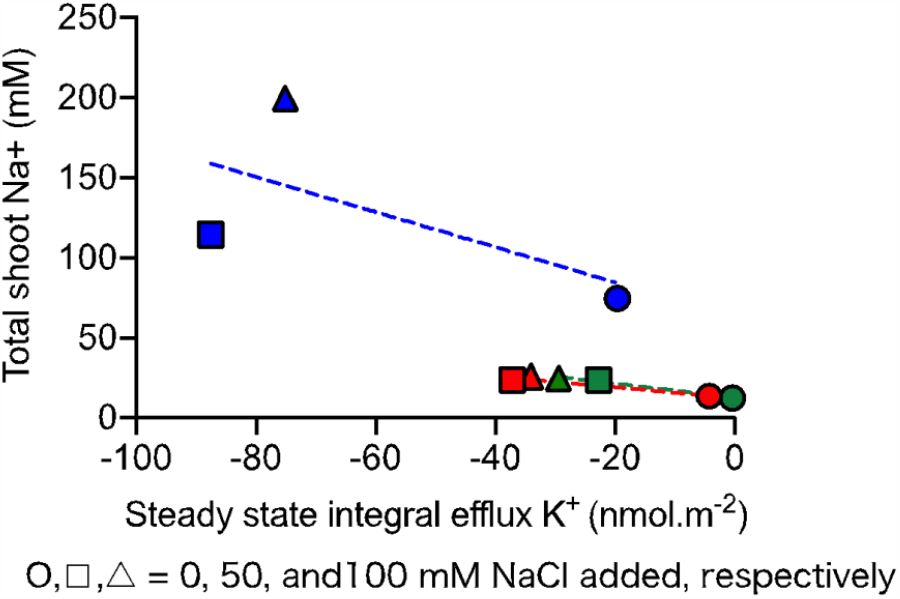
The relation between the integrated K^+^ efflux characteristics and the total Na^+^ accumulated in the shoot of the three grass species. Green symbols represent the values for *Puccinellia maritima*, red for *Festuca rubra*, and blue for *Lolium perenne*. The data from Figure 2 (K^+^ flux data) and Figure 4 (contribution of NaCl to total osmolarity) were used to calculate the total K^+^ efflux (summation ‘flux x time’ of every timepoint over the measuring period) and the relative concentration of Na^+^ in the shoot. In the figure it is evident that there is a clear correlation between these fluxes and that there is a clear difference between on the one hand *F. rubra* and *P. maritima* and on the other *L. perenne*.

**Figure 7.**
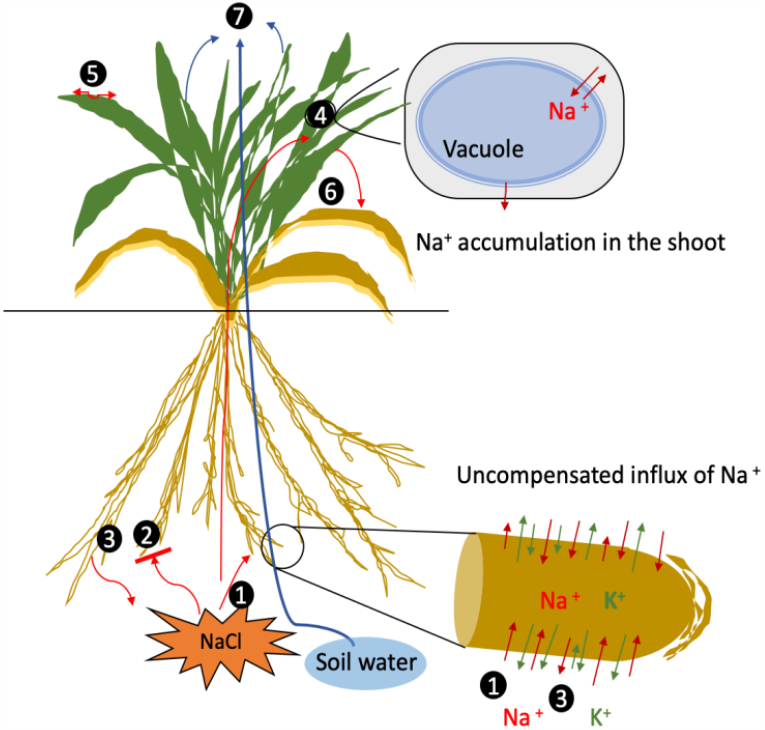
Schematic representation of the different processes involved in the regulation of Na^+^ transport and accumulation in the shoot where it can cause damage to cellular processes when the K^+^/Na^+^ ratio is lowered or contribute to the overall cell osmotic adjustment when compartmentalized in the vacuole. ➀. Na^+^ uptake in the root cells;➁. NaCl exclusion from the root; ➂.NaCl efflux from root cells; ➃. NaCl long-distance transported to the leaf and stored in the vacuole; ➄.NaCl exclusion from leaf after long-distance transport from the root; ➅.NaCl reallocation to older leaves; ➆. Water evaporation.

### 4.2. K^+^ fluxes recovery capacity as an indicator of salt tolerance

The differences in root H^+^ and K^+^ flux responses to salinity among the three tested grass species are correlate with their variations in salinity tolerance. The limited Na^+^-triggered K^+^ efflux, and net K^+^ inward fluxes, coupled with the activity of the H^+^ pump, contribute to homeostasis of the cell K^+^/Na^+^ ratio. Cytosolic K^+^/Na^+^ ratio has been identified as a critical determinant of plant salt tolerance(Shabala & Pottosin, 2014; Assaha *et al.*, 2017). The physicochemical similarity of Na^+^ and K^+^ leads to competition for essential binding sites on enzymes, where only K^+^ can render the enzyme functional (Assaha et al., 2017). When Na^+^ is abundant, it can disrupt physiological processes by displacing K^+^ from these binding sites. Therefore, it is crucial to limit K^+^ loss and prevent Na^+^ from entering the root. Halophytic species, which thrive in high salinity conditions, typically possess superior mechanisms for preventing K^+^ leakage(Flowers & Colmer, 2008; Assaha *et al.*, 2017).

The reversal of Na^+^-triggered K^+^ efflux to a K^+^ influx relies on the repolarization of the membrane driven by H^+^ efflux through plasma membrane-bound ATPases. The three species studied varied considerably in capacity to exhibited considerable variation in their ability to limit K^+^ and H^+^ efflux when exposed to Na^+^. Both *L. perenne* and *F. rubra* displayed a substantial transient K^+^ efflux in response to Na^+^, whereas *P. maritima* showed absent K^+^ efflux. In comparison to K^+^ fluxes, the net H^+^ flux was much lower, ranging from 0 to less than -200 nmol·m-2·s-1. Additionally, *P. maritima* exhibited the lowest H^+^ fluxes among the three species, whereas *F. rubra* showed a significant Na^+^-induced K^+^ efflux. However, the transient reverse of the influx of H^+^ might indicate a subsequent recovery of the membrane potential and a driving force for K^+^ influx. *L. perenne* displayed the lowest ability in both the retention of K^+^ and the recovery of K^+^ efflux.

### 4.3. Osmolarity and NaCl contribution as the measure of salt tolerance

Salinity exerts detrimental effects on plants by imposing both osmotic and ionic stress. The high external osmolarity caused by salt can lead to water loss from plant cells, resulting in reduced cell turgor and slowed elongation. Halophytes and glycophytes differ in their ability to maintain turgor under high salinity conditions(Hasegawa *et al.*, 2000; Munns & Tester, 2008). The osmosensor, as shown in Figure 5, activates H^+^-ATPase transporters at the cell membrane, leading to hyperpolarization of the membrane potential and the proton motive force. These forces can be utilized to drive secondary ion transport across the membrane (Gaxiola *et al.*, 2007).

Plants have two main ways to restore osmotic balance between the cytoplasm and the external medium. One way is through the direct uptake of available ions such as Na^+^, K^+^, and Ca2^+^. The other method involves the synthesis of compatible solutes, which are organic compounds that help maintain osmotic balance(Li *et al.*, 2009). Roots play a crucial role in these processes as they are involved in water uptake, ion uptake, and regulation, as well as the relocation of ions within the plant. In the case of *L. perenne*, the loss of K^+^ and the high contribution of NaCl to the total osmolarity in leaf sap indicate that NaCl has entered the plant root and been transported to the shoot, where it significantly contributes to the total osmolarity under salinity.

While many true halophytes are known for their ability to take up Na^+^ as a cheap inorganic osmoticum and store it in the vacuole, *P. maritima* mihgt follow a different mechanism. It synthesizes organic solutes to maintain osmotic balance(Fokkema *et al.*, 2016). *P. maritima* might restricts the leakage of K^+^ from the root, effectively excludes Na^+^ compared to *L. perenne* and *F. rubra*, and maintains an adequate osmotic balance, with only a limited contribution of Na^+^ in the shoot. These osmotically active compounds synthesized by *P. maritima*, such as amino acids, amides, proteins, quaternary ammonium compounds, and polyamines(Hasegawa *et al.*, 2000; Munns & Tester, 2008), explain why it is favored by small herbivores and birds in the salt marsh, selecting it for its high nutritious value(Fokkema *et al.*, 2016).

## Acknowledgments

We would like to thank the students Wycher Wind and Niels Helfferich for their photos of the root cross sections. J. Yi is thanked for editing the references and SM as required. **Funding:** LW was funded by the Chinese Scholarship Council: CSC number 201600090216

## Author Contributions

MS, LW, and TE designed the experiments and wrote the manuscript; LW and MS performed the experiments and the data collection; LW, TE, and MS did the data analyses. All authors have read and agreed to the published version of the manuscript.

## Data Availability Statement

The data that support the findings of this study are openly available in https://dataverse.nl/dataverse/GELIFES

## Supplementary Materials

The following supporting information can be downloaded at: https://figshare.com/account/projects/175446/articles/23937768. Box S1: Material of microelectrodes and preparation, Box S2: Making selective microelectrodes and calibration. Figure S1: The measurement of ion flux and two distant point of root surface. Figure S2: Effect of salinity on the accumulation of dry matter of in the shoot of *Lolium perenne, Festuca rubra* and *Puccinellia maritima*. Table S1: peak fluxes location along root in control and salt condition in three grass species. Table S2: P-value details of Leaf sap osmolarity and contribution of NaCl to the total osmolarity of plant leaf sap. Table S3: Two-way Anova for Total dry matter in the shoot of *Lolium perenne, Festuca rubra* and *Puccinellia maritima*.

